## Supplementary Tables and Figures for "Design, 3D-printing, and characterisation of a low-cost, open-source centrifuge adaptor for separating large volume clinical blood samples"

**Supplementary Table 1:** Technical specification of SciSpin MINI Microfuge, model: SQ-6050

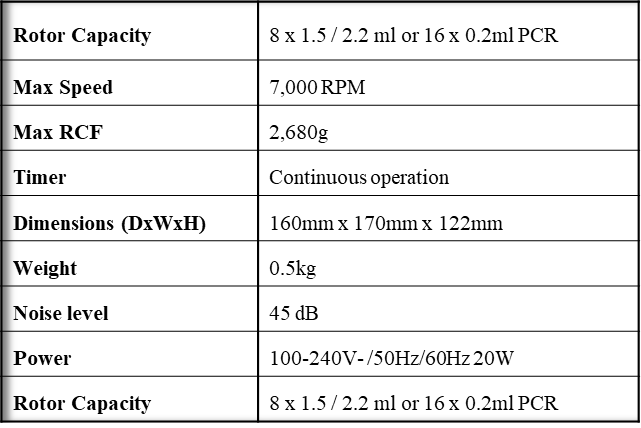

**Supplementary Table 2:** Technical characteristics of Anycubic i3 Mega 3D printer

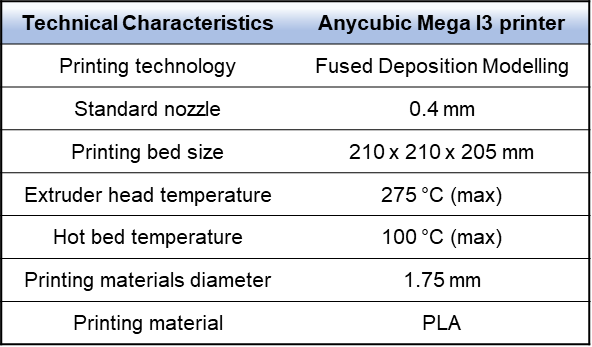

**Supplementary Table 3:** 3D printing setting for Ultimaker Cura 4.4

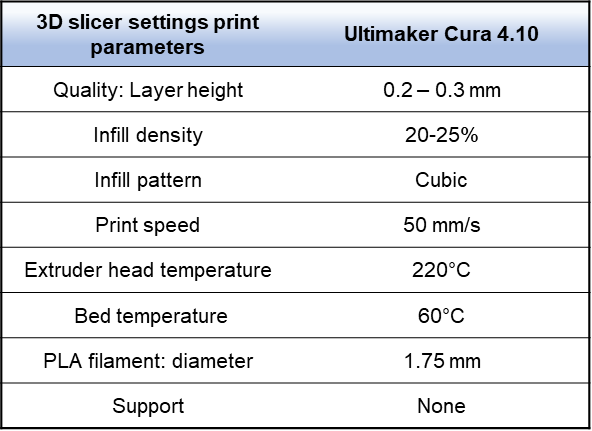

**Supplementary Table 4:** Motor power load of different designs at the loaded and unloaded condition

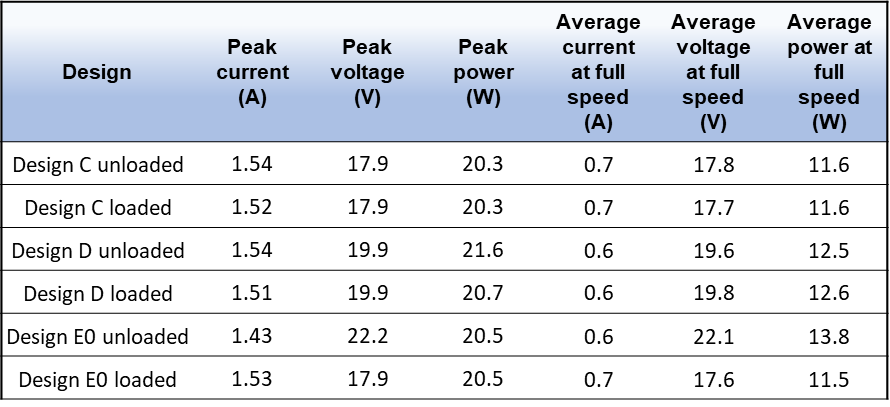

**Supplementary Figure S1**

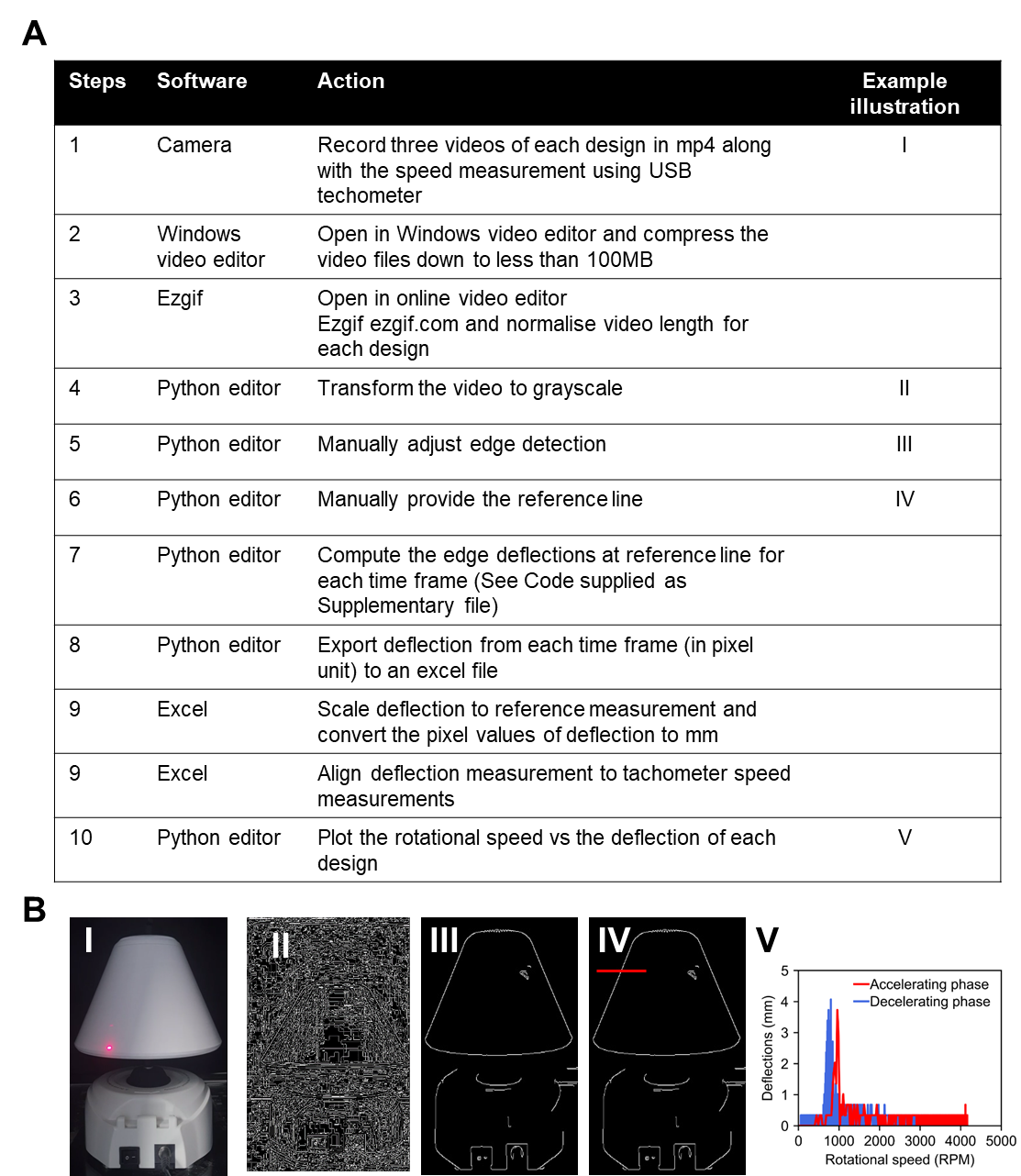

**Supplementary Figure S1:** **(A)** Step-by-step guide for the deflection measurements **(B)** Corresponding illustration for steps annotated I-V in (A). One video of design C has been uploaded as a separate Supplementary File as an example. All three video files for each design are available from FigShare https://doi.org/10.6084/m9.figshare.16762444.v1

**Supplementary Figure S2**

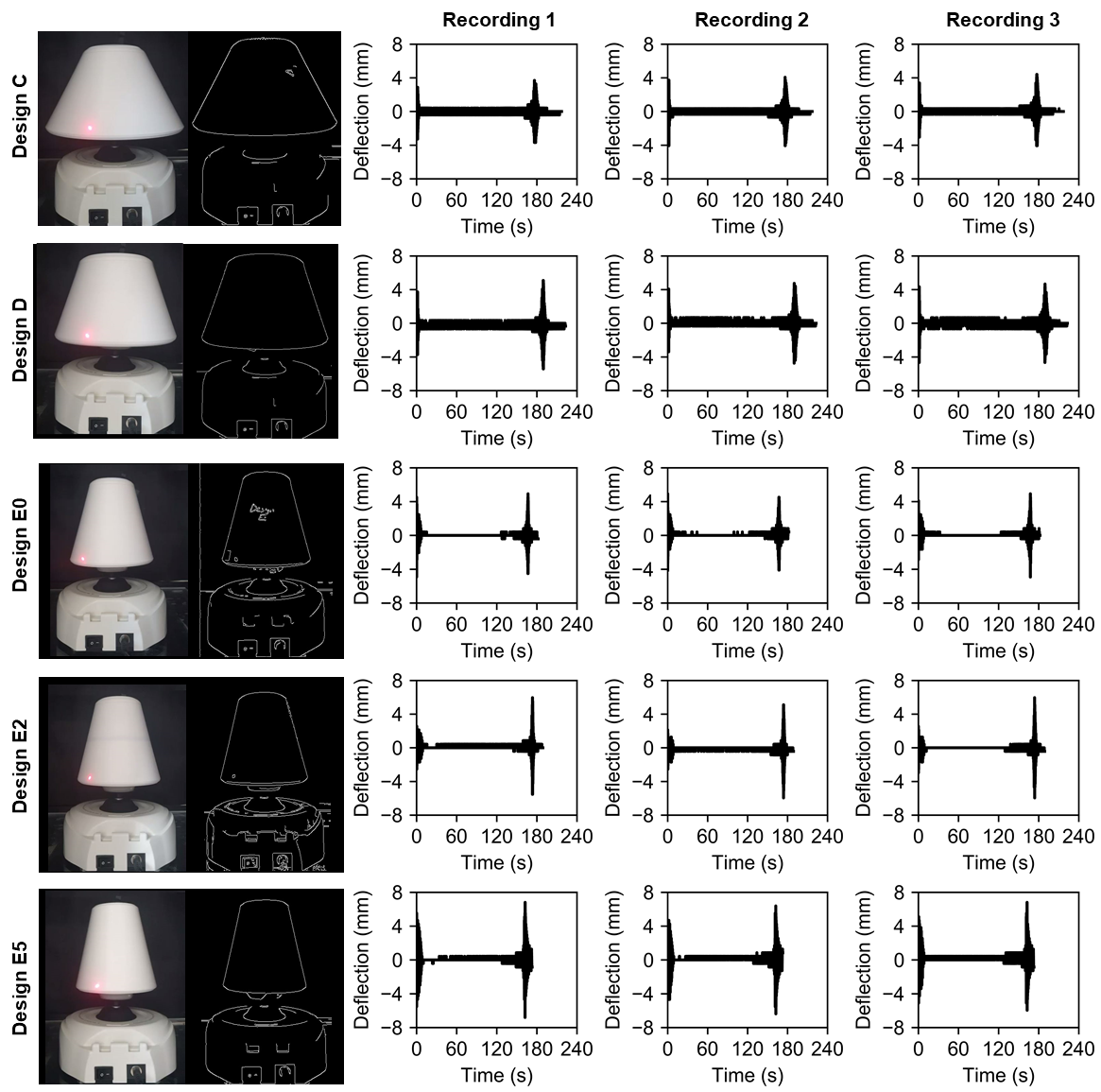

**Supplementary Figure S2:** Real-time deflection measurement from the recorded video of Designs C, D, E0, E2, E5

**Supplementary Figure S3**

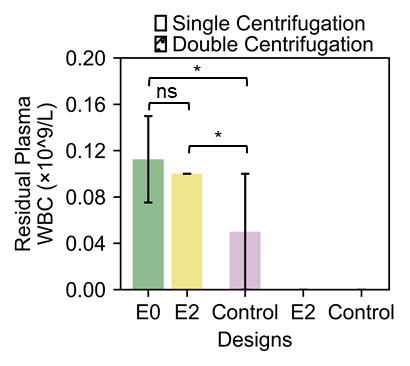

**Supplementary Figure S3**: WBC count on the residual plasma after single centrifugation and double centrifugation (A second centrifugation @12000 RCF, 10 minutes)
