## Supplementary material for "Design, 3D-printing, and characterisation of a low-cost, open-source centrifuge adaptor for separating large volume clinical blood samples": Python code for vibration measurements

Python code for deflection measurement from recorded video

**#Matplotlib inline library**

import cv2

import numpy as np

import time

import matplotlib.pyplot as plt

import os

import pandas as pd

from openpyxl import load_workbook

cv2.namedWindow("window")

def nothing(x):

pass

**# for finding optimal values**

cv2.createTrackbar("upper", "window", 0, 255, nothing)

cv2.createTrackbar("lower", "window", 0, 255, nothing)

vid = cv2.VideoCapture('Documents/matplotlib/3rd phase/corrected/c2.mp4')

cap = cv2.VideoCapture('Documents/matplotlib/3rd phase/corrected/c2.mp4')

_, img = cap.read()

img = cv2.resize(img, (450, 700))

gray = cv2.cvtColor(img, cv2.COLOR_BGR2GRAY)

while True:

**# sets x and y which will be used later for edge detection**

x = cv2.getTrackbarPos('lower', 'window')

y = cv2.getTrackbarPos('upper', 'window')

edge = cv2.Canny(gray, x, y)

cv2.imshow('window', edge)

if cv2.waitKey(1) == 27:

cv2.destroyAllWindows()

break

orig = 76

diff = []

initial = False

while True:

try:

_, frm = vid.read()

frm = cv2.resize(frm, (450, 700))

gray = cv2.cvtColor(frm, cv2.COLOR_BGR2GRAY)

img = cv2.Canny(gray, x, y)

**#saving one image for pixel calculation**

path = 'Documents/matplotlib/3rd phase/corrected/'

cv2.imwrite(os.path.join(path , 'c2.jpg'), img)

**# thresholding to only have either pixel value 0 or 255**

_, img = cv2.threshold(img, 0, 255, cv2.THRESH_BINARY)

cv2.imshow("winname", img)

**# refrence line**

gr = img[220, 115:170]

#gr = img[233, 30:70]=255

#cv2.imshow("gr", img)

if not(initial):

**# setting refrence point**

ref = np.argmax(gr)

initial = True

**# getting current positon**

curr_val = np.argmax(gr)

print(gr)

print(ref)

print(curr_val)

print(ref-curr_val)

**# getting and appending deflection from ref point**

if curr_val != 0:

diff.append(ref-curr_val)

else:

diff.append(0)

except:

break

if cv2.waitKey(1) == 27:

break

cv2.destroyAllWindows()

print(diff)

t = np.arange(len(diff))

**#writing to specific excel sheet using path & df to excel**

path= r"Documents/matplotlib/3rd phase/corrected/3rd phase.xlsx"

book = load_workbook(path)

writer = pd.ExcelWriter(path, engine = 'openpyxl')

writer.book = book

#writer = pd.ExcelWriter(path, engine = 'xlsxwriter')

df = pd.DataFrame(diff)

df.to_excel(writer, sheet_name = 'C2_XY')

writer.save()

writer.close()

\# plotting everything

plt.plot(t ,diff)

plt.show()
